# Human cardiomyocytes with trisomy 21 exhibit heightened susceptibility and immunological response to SARS-CoV-2 infection

**DOI:** 10.64898/2026.06.22.733265

**Authors:** Matthew Alonzo, KC Mahesh, Cankun Wang, Jerry Wang, Jakob Bering, Qin Ma, Vidu Garg, Mark Peeples, Ming-Tao Zhao

## Abstract

Coronavirus disease 2019 (COVID-19), caused by SARS-CoV-2, is associated with significant cardiovascular complications, including myocardial injury and long-term cardiac dysfunction. Individuals with Down syndrome (trisomy 21) exhibit increased susceptibility to severe COVID-19 outcomes, yet the cardiomyocyte-intrinsic mechanisms underlying this vulnerability remain poorly understood. To investigate genotype-specific responses to SARS-CoV-2 infection, we generated induced pluripotent stem cell–derived cardiomyocytes from individuals with trisomy 21 and their euploid, sex-matched biological relatives. Cardiomyocytes were inoculated with SARS-CoV-2, and viral susceptibility was assessed by immunofluorescence. Bulk RNA sequencing was performed under baseline and infected conditions to define transcriptional programs associated with viral response. Trisomy 21 iPSC-CMs exhibited increased susceptibility to SARS-CoV-2 infection, with greater viral protein expression and a higher proportion of infected cardiomyocytes compared to controls. Baseline transcriptomic analysis revealed no significant differences in canonical viral entry factors including *ACE2* and *TMPRSS2*, suggesting that differential susceptibility is not driven by entry receptor availability. Following infection, both trisomy 21 and euploid control groups activated conserved antiviral pathways; however, trisomy 21 cardiomyocytes displayed a markedly amplified transcriptional response, with substantially greater numbers of differentially expressed genes. Upregulated pathways included interferon signaling, NF-κB activation, cytokine and chemokine signaling, and innate immune responses, while downregulated pathways were enriched for cardiomyocyte structural integrity, calcium handling, and metabolic processes. Notably, inflammatory and cytokine-related transcripts were significantly more elevated in trisomy 21 cells, consistent with an exaggerated immune response. These findings provide mechanistic insight into the increased cardiovascular risk observed in individuals with Down syndrome and highlight dysregulated immune signaling as a potential therapeutic target in this high-risk population.

## 1. Introduction

Coronavirus disease 2019 (COVID-19), caused by severe acute respiratory syndrome coronavirus 2 (SARS-CoV-2), is recognized as a multisystem disorder with significant cardiovascular involvement. Myocardial injury occurs in approximately 20–30 % of hospitalized patients and is associated with adverse outcomes, including arrhythmias, myocarditis, and increased mortality [1]. Persistent cardiac abnormalities in convalescent individuals further suggest that SARS-CoV-2 can induce long-term cardiovascular dysfunction.

These clinical observations point to complex mechanisms of cardiac injury involving both direct and indirect effects of viral infection. Inflammation, hypoxia, and microvascular dysfunction contribute to tissue damage, while direct infection of cardiomyocytes is increasingly supported by evidence demonstrating expression of angiotensin-converting enzyme 2 (ACE2), the primary viral entry receptor, and *in vitro* studies showing permissiveness of human induced pluripotent stem cell-derived cardiomyocytes (iPSC-CMs) to SARS-CoV-2 infection [2–4]. Detection of viral RNA and particles in human cardiac tissue further substantiates this mechanism. However, because of its original characterization as primarily a respiratory disease, mechanistic insight into cardiomyocyte-specific responses to SARS-CoV-2 infection remains limited.

Understanding cardiomyocyte-specific response to SARS-CoV-2 infection is particularly relevant in populations with heightened vulnerability to SARS-CoV-2. Individuals with Down syndrome (DS), or trisomy 21, represent one such high-risk group. Affecting approximately 1 in 800 live births, DS is characterized by developmental delays, immune dysregulation, and a high prevalence (∼50 %) of congenital heart defects [5]. Epidemiological studies consistently report increased hospitalization and mortality following SARS-CoV-2 infection in this population, yet the biological basis for the increased susceptibility remains poorly defined [6–8]. Although the acute phase of the COVID pandemic has subsided, SARS-CoV-2 continues to circulate and contributes to chronic health conditions, including long COVID and persistent cardiovascular complications [9]. Elucidating the mechanisms of cardiac injury, particularly in vulnerable populations such as individuals with Down syndrome, remains essential for improving long-term clinical management, guiding future therapeutic strategies, and enhancing preparedness for emerging viral pathogens with similar cardiovascular impacts.

In the present study, we sought to define the cardiomyocyte-specific mechanisms underlying SARS-CoV-2–induced pathology in DS. To address this, we utilized iPSC-CMs from individuals with trisomy 21 and euploid, sex-matched biological relatives, enabling the investigation of genotype-specific responses. By combining viral infection assays with transcriptomic profiling, we aimed to characterize how trisomy 21 influences susceptibility to SARS-CoV-2 infection and the associated cellular response programs. These studies provide a framework for understanding the molecular basis of increased COVID severity in individuals with DS.

## 2. Results

We differentiated human iPSCs from three unique trisomy 21 subjects and each subject’s euploid sex-matched biological parent into cardiomyocytes using published protocols that involves modulating Wnt signaling [10]. Resultant cardiomyocytes beat spontaneously in culture and showed intercalated α-ACTININ and cardiac TNNT2 expression (**Figure 1A**). Euploid iPSC-CMs from matched controls showed organized and robust sarcomere structures compared to disordered and diffuse contractile proteins observed in cells with trisomy 21.

**Figure 1.**
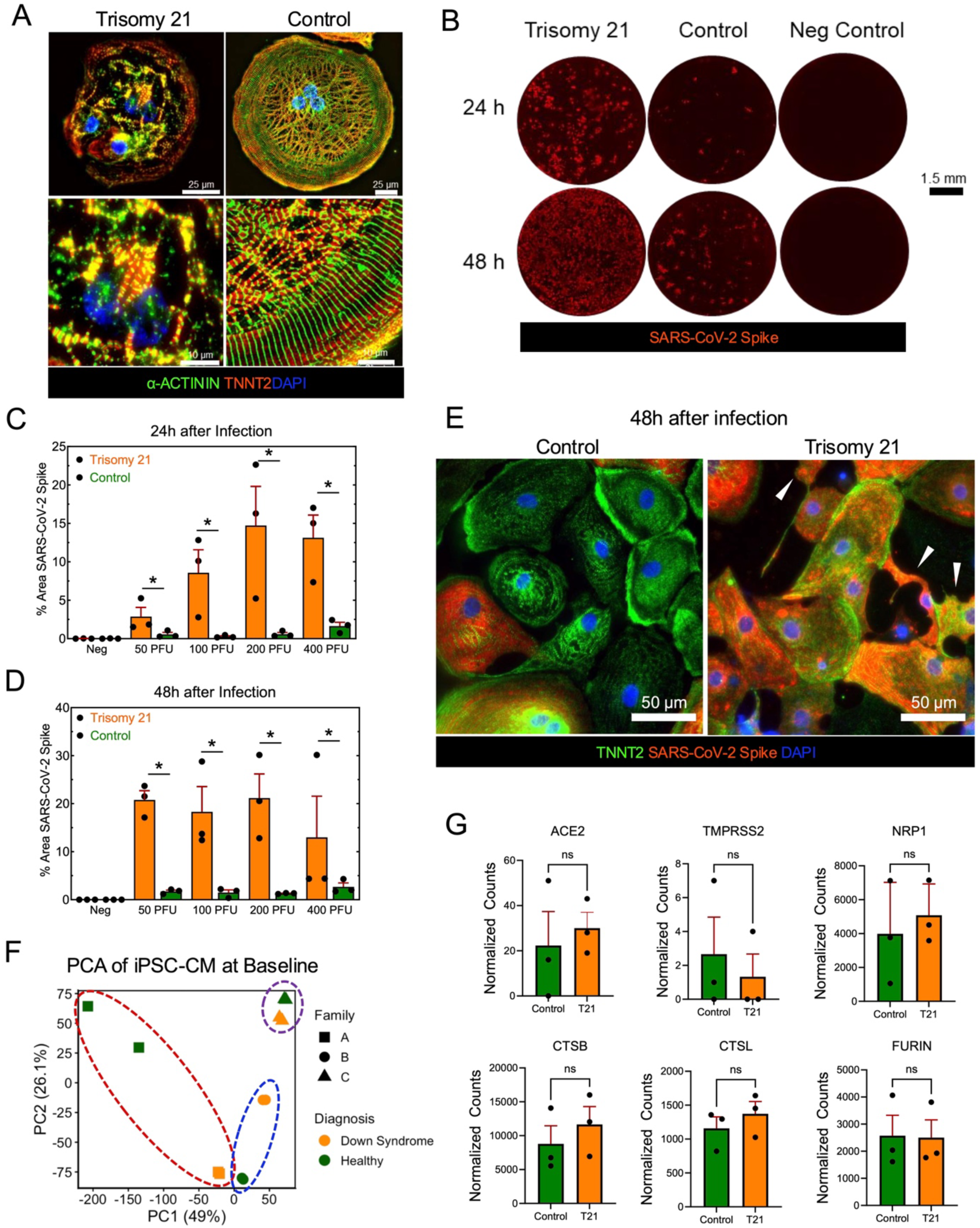
Trisomy 21 iPSC-CMs exhibit increased susceptibility to SARS-CoV-2 infection. **(A)** Control iPSC-CMs displayed organized sarcomeres (TNNT: green, α-ACTININ: red), whereas trisomy 21 iPSC-CMs showed disrupted architecture. Nuclei were stained with DAPI (blue). Scale bars: 25 μm (top), 10 μm (bottom). **(B)** At 24 h and 48 h post-infection, trisomy 21 iPSC-CMs demonstrated higher SARS-CoV-2 spike protein expression (red) and greater viral spread relative to controls. Scale bar: 1.5 mm. **(C–D)** Quantification of spike protein fluorescence confirmed significantly elevated signal at 24 h (C) and enhanced spread at 48 h (D) in trisomy 21 iPSC-CMs (*p < 0.05, paired Student’s t-test). **(E)** High-magnification immunofluorescence images of control and trisomy 21 iPSC-CMs at 48 h post–SARS-CoV-2 infection. In control cultures, TNNT2⁺ cardiomyocytes (green) exhibit heterogeneous spike protein expression (red), with a subset of infected cells. In contrast, trisomy 21 cultures display a greater proportion of spike-positive cells, accompanied by pronounced abnormalities in cardiomyocyte morphology highlighted by arrowheads. Nuclei are stained with DAPI (blue). Scale bar: 50 μm. **(F)** PCA of baseline transcriptomic profiles from trisomy 21 and euploid iPSC-CMs demonstrates clustering by familial pairs along PC1 (circled), with separation by karyotypes observed within each pair. (**G**) Expression of selected transcripts involved in SARS-CoV-2 host cell entry (*ACE2, TMPRSS2, NRP1, CTSB, CTSL, FURIN*) shows no significant differences between trisomy 21 and control uninfected iPSC-CMs. Wilcoxon matched-pairs signed-rank test. *ACE2 p* = 0.75, *CTSB p* = 0.75, *CTSL p* = 0.25, *FURIN p* > 0.999, *NRP1 p* = 0.25.

DS and Control cardiomyocytes were matured to day 30 and then inoculated with various doses (0, 50, 100, 200, 400 PFU) of SARS-CoV-2 in a 96-well plate to compare viral infectivity. After 24h and 48h of infection, the SARS-CoV-2 spike protein was detected via immunofluorescence. At 24 and 48 h post-infection, the coronavirus spike protein fluorescence signal encompassed a larger area in trisomy 21 samples compared to controls (**Figure 1B**). We quantified the area of fluorescence for each viral dosage and found significantly higher levels of spike protein-expressing iPSC-CMs in trisomy 21 for all viral dosages at 24 and 48 h compared to matched controls (**Figure 1C-1D**). Magnified immunofluorescence images of TNNT2⁺ cardiomyocytes demonstrated that infection was limited to a subset of control cardiomyocytes, with little to no detectable coronavirus spike protein in the majority of cells. In contrast, trisomy 21 cardiomyocytes exhibited widespread infection, with the majority of iPSC-CMs showing robust spike protein expression. These results indicate increased DS cardiomyocyte susceptibility to SARS-CoV-2 infection and an ability of the virus to spread more rapidly in DS cells than in euploid control cells (**Figure 1E**). Furthermore, a subset of infected cardiomyocytes in the DS group exhibited pronounced morphological abnormalities (highlighted by white arrowheads), indicating a more severe and distinct cytopathic response to infection. Given these pronounced differences in viral burden and cellular phenotype, we next sought to understand the underlying transcriptional landscape of control and DS iPSC-CMs at baseline (absent SARS-CoV-2 infection) and in response to infection.

RNA was isolated from uninoculated day 30 iPSC-CMs and 48 h after SARS-CoV-2 infection and subjected to bulk RNA sequencing to generate global transcriptional profiles for each group. Principal component analysis (PCA) of baseline transcriptomic data revealed that the greatest proportion of variance (PC1) was attributable to genetic background, as samples clustered primarily by family origin (**Figure 1F**). Notably, within each family-specific cluster, a clear segregation between trisomy 21 and euploid control samples was observed, indicating that disease status contributes an additional layer of transcriptional variability.

At baseline, analysis of genes implicated in SARS-CoV-2 cellular entry did not reveal any statistically significant differences between trisomy 21 and control iPSC-CMs (**Figure 1G**). Expression of *ACE2*, the principal receptor mediating SARS-CoV-2 attachment [11], was detectable in both groups. In contrast, transcripts encoding transmembrane serine protease 2 (*TMPRSS2*), a key protease responsible for coronavirus spike protein priming at the cell surface [12] were minimally expressed or absent in both DS and control groups. The near absence of *TMPRSS2* transcripts suggests canonical plasma membrane fusion is unlikely to be the predominant route of viral entry in these cells, suggesting alternative endocytic mechanisms. Consistent with this, both trisomy 21 and control iPSC-CMs exhibited enrichment of lysosomal cysteine proteases cathepsin B (*CTSB*) and cathepsin L (*CTSL*), which are known to facilitate SARS-CoV-2 entry via the endosomal pathway through proteolytically activating of the viral spike protein [13]. Additional host factors associated with viral uptake, including furin (*FURIN*) and neuropilin-1 (*NRP1*), were also enriched in both trisomy 21 and control cardiomyocytes at baseline.

Given that baseline differences in viral entry factors did not account for the increased susceptibility observed in trisomy 21 cells, we next examined the transcriptional response to SARS-CoV-2 infection by performing differential expression analyses comparing infected and baseline conditions within each group. PCA of post-infection samples again demonstrated that the largest source of variance (PC1) was driven by genetic background, with RNA samples clustering by family. Within each family cluster, separation by diagnosis remained evident, indicating that both host genetics and trisomy 21 status contribute to the transcriptional response to infection (**Figure 2A**).

**Figure 2.**
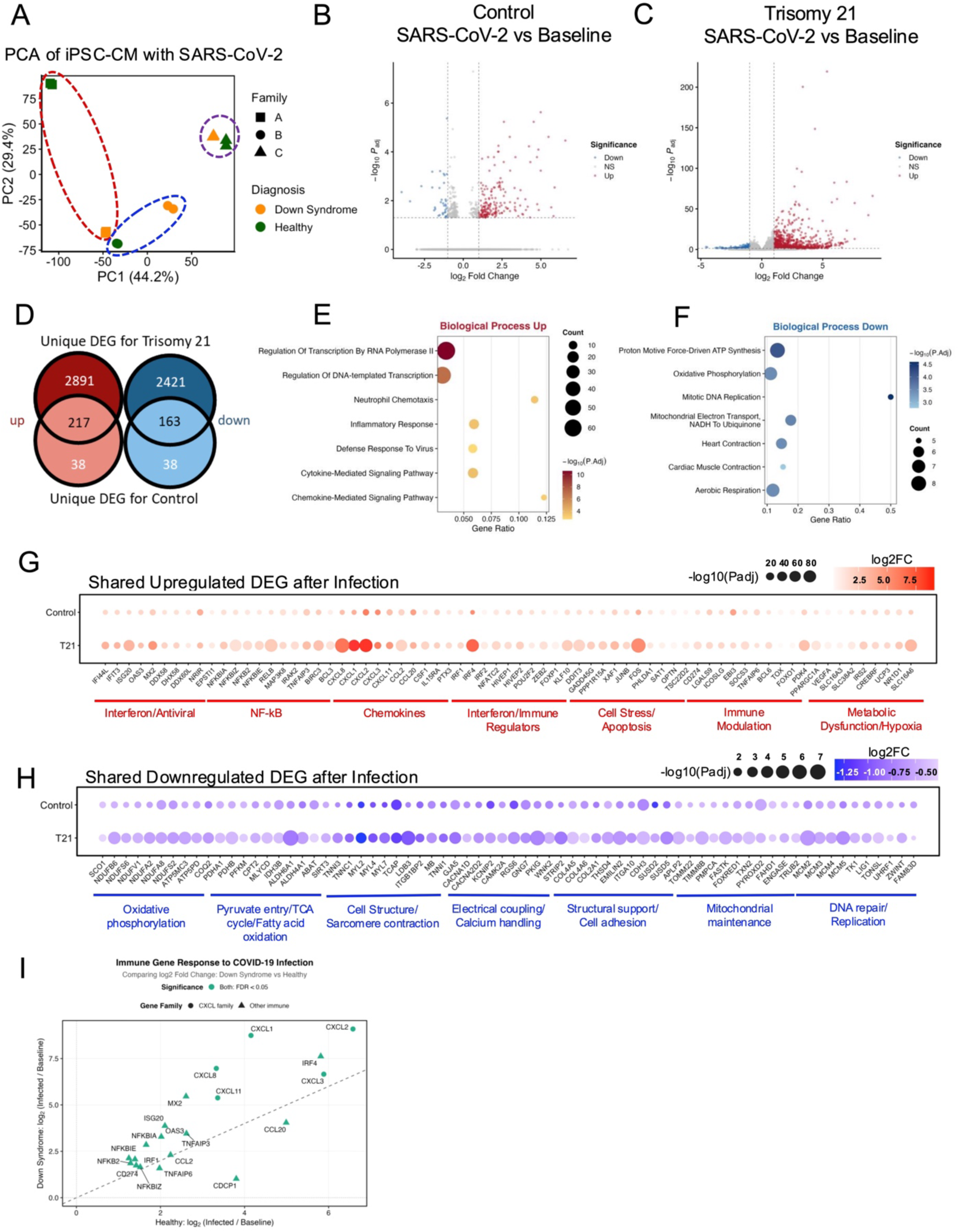
Transcriptomic signatures of viral infection in trisomy 21 and euploid cardiomyocytes show amplified inflammatory response in Down syndrome. **(A)** PCA plot of global transcriptome analysis of trisomy 21 and control iPSC-CMs after SARS-CoV-2 infection. Datapoints separated by familial pairs along PC1 (circled pairs) with separation within each family by karyotype. (**B–C**) Volcano plots of DEG following SARS-CoV-2 infection in (B) euploid controls and (C) trisomy 21 samples (threshold: Log2FC ≥ 2, Padj ≤ 0.05). (**D**) Venn diagram illustrates the number of uniquely upregulated and downregulated transcripts in trisomy 21 and control groups after SARS-CoV-2 infection, as well as overlapping transcripts shared between groups after viral infection. (**E–F**) Gene ontology (GO) enrichment analysis for (E) shared upregulated biological processes and (F) shared downregulated transcripts between groups. (**G**) Representative shared upregulated (red) transcripts are enriched for pathways related to inflammation, antiviral response, cellular stress, and apoptosis. (**H**) Representative shared downregulated (blue) transcripts are enriched for processes related to metabolism, sarcomere function, calcium handling, and DNA damage/replication indicating impaired cardiomyocyte homeostasis following infection. (**I**) Comparative analysis of immune-related transcript enrichment between trisomy 21 (y-axis) and control (x-axis) samples demonstrate increased enrichment in trisomy 21 as indicated by most transcripts falling above the diagonal. Transcripts along the diagonal exhibit comparable expression between groups, whereas transcripts below the diagonal are enriched in control samples and those above are enriched in trisomy 21.

Differential gene expression analysis revealed striking differences in the magnitude of the transcriptional response between groups. Volcano plots comparing infected samples to their respective baselines showed a markedly amplified transcriptional response in trisomy 21 iPSC-CMs (**Figure 2C**) relative to controls (**Figure 2B**). Specifically, trisomy 21 samples exhibited 2,891 unique upregulated and 2,421 unique downregulated transcripts, whereas cells from matched controls demonstrated a comparatively modest response, with only 38 unique upregulated and 38 unique downregulated transcripts (**Figure 2D**). In addition, 217 transcripts were upregulated in common, and 163 transcripts were downregulated in common across groups, representing a conserved core response to SARS-CoV-2 infection.

Gene ontology and pathway enrichment analysis of the shared upregulated transcripts revealed activation of antiviral and inflammatory pathways, including cytokine- and chemokine-mediated signaling pathways, and innate immune responses (**Figure 2E**). These findings are consistent with viral infection and a coordinated host response aimed at restricting viral replication and promoting immune activation. Conversely, shared downregulated transcripts were enriched for pathways critical to cardiomyocyte function, including sarcomere organization and cardiac muscle contraction, calcium ion handling (e.g., excitation–contraction coupling), mitochondrial energetics, and oxidative metabolism (**Figure 2F**). Suppression of these pathways suggests that SARS-CoV-2 infection induces a shift away from normal contractile and metabolic functions toward a stress- and immune-responsive state, potentially contributing to cardiomyocyte dysfunction.

Visualization of selected shared upregulated genes using a bubble plot further highlighted increased expression of interferon-stimulated genes, NF-κB target genes, pro-inflammatory cytokines and chemokines (e.g., CXCL and CCL family members), as well as regulators of innate immunity and apoptosis (**Figure 2G**). These transcriptional changes reflect robust activation of antiviral defense mechanisms and inflammatory signaling cascades. In contrast, the downregulated gene set (**Figure 2H**) included key regulators of cardiac contractility (e.g., structural and sarcomeric proteins), calcium cycling proteins, and metabolic enzymes involved in mitochondrial respiration and ATP production, reinforcing the notion of functional impairment in infected cardiomyocytes.

Given that the well-documented increased severity of COVID outcomes in individuals with DS is often attributed to the cytokine storm following infection, we directly compared inflammatory and cytokine-related gene expression between trisomy 21 and control iPSC-CMs. Analysis of log2 fold changes (infected versus baseline) revealed a pronounced enrichment of pro-inflammatory cytokines, chemokines, and immune signaling mediators in trisomy 21 iPSC-CMs relative to controls (**Figure 2I**). This included heightened expression of genes associated with cytokine storm phenotypes such as interferons, interleukins, and chemokine ligands, indicating an exaggerated and potentially maladaptive antiviral response.

## 3. Discussion

In the present study, we investigated the cardiomyocyte-intrinsic response to SARS-CoV-2 infection using iPSC-derived cardiomyocytes from individuals with trisomy 21 and euploid, genetically related sex-matched parental controls. Our findings demonstrate that trisomy 21 cardiomyocytes exhibit increased susceptibility to viral infection and mount a markedly increased transcriptional response characterized by heightened inflammatory signaling and suppression of core cardiomyocyte functional programs. These results may provide mechanistic insight into the increased morbidity and mortality observed in individuals with DS following SARS-CoV-2 infection.

A key observation from our study is the significantly greater extent of infection in trisomy 21 iPSC-CMs, as evidenced by increased spike protein signal intensity and a broader distribution of infected TNNT2⁺ cardiomyocytes. This enhanced susceptibility was not explained by baseline differences in canonical viral entry transcripts, as *ACE2* expression was comparable between groups and *TMPRSS2* expression was minimal in both conditions. Instead, our data suggest that SARS-CoV-2 entry into iPSC-CMs likely occurs through alternative, endosome-mediated pathways involving cathepsins (*CTSB*, *CTSL*), *FURIN*, and *NRP1*. Yet, similar expression levels of these factors between groups indicate that differential viral entry alone is unlikely to account for the increased infection burden in trisomy 21 cells.

Furin-mediated cleavage of the spike protein enhances its conformational readiness for receptor engagement, while *NRP1* has been shown to augment viral infectivity by promoting viral internalization. The concurrent expression of these transcripts supports the possibility of parallel or functionally redundant entry pathways operating in iPSC-CMs. Collectively, these findings suggest that SARS-CoV-2 entry in iPSC-CMs is likely mediated through non-canonical, endosome-associated mechanisms rather than *TMPRSS2*-dependent plasma membrane fusion. However, despite comparable expression of key entry factors between trisomy 21 and control cells, these baseline transcriptomic data do not fully explain the increased susceptibility to SARS-CoV-2 infection observed in DS-derived iPSC-CMs. Additional factors, potentially involving post-transcriptional regulation, host antiviral responses, or differences in cellular physiology may contribute to the heightened vulnerability associated with trisomy 21.

The most striking difference between trisomy 21 and control cardiomyocytes emerged at the level of the transcriptional response to SARS-CoV-2 infection. While both groups activated a conserved antiviral program including interferon signaling, NF-κB activation, and cytokine-mediated pathways, trisomy 21 cells exhibited a dramatically heightened response, with orders-of-magnitude greater numbers of differentially expressed genes. This hyperactivation of immune and inflammatory pathways is consistent with prior reports of immune dysregulation in Down syndrome and may reflect gene dosage effects arising from chromosome 21 [14]. Notably, chromosome 21 encodes several interferon receptor subunits, which have been implicated in hypersensitivity to interferon signaling [15]. Overexpression of these receptors could prime trisomy 21 cells for an exaggerated response to viral infection, resulting in enhanced induction of interferon-stimulated genes and downstream inflammatory cascades.

The pronounced enrichment of pro-inflammatory cytokine and chemokine transcripts in trisomy 21 iPSC-CMs is consistent with a model in which dysregulated immune signaling contributes to disease severity. Elevated expression of chemokine and cytokine transcripts may not only promote local inflammation within the myocardium, but also amplify systemic immune responses, potentially contributing to the “cytokine storm” phenotype observed in severe COVID cases. In the context of DS, where baseline immune homeostasis is already perturbed, this exaggerated response may be particularly detrimental, leading to increased tissue damage and impaired recovery.

Despite these insights, several limitations should be considered. Human iPSC-CMs, while a powerful model system, do not fully recapitulate the maturity and multicellular complexity of adult human myocardium. The absence of immune cells, fibroblasts, and endothelial components limits our ability to model cell–cell interactions that are critical *in vivo*. Additionally, bulk RNA sequencing captures averaged transcriptional responses and may obscure cell-to-cell heterogeneity in infection and response. Future studies employing single-cell transcriptomics, proteomics, and functional assays will be important to further dissect these mechanisms. Moreover, investigation into post-transcriptional regulation, including protein expression, signaling dynamics, and viral replication kinetics, will be necessary to fully understand the basis of increased susceptibility in trisomy 21 cardiomyocytes.

In summary, our study demonstrates that trisomy 21 cardiomyocytes are intrinsically more susceptible to SARS-CoV-2 infection and exhibit a heightened inflammatory transcriptional response compared to euploid sex-matched controls. These findings provide a mechanistic framework linking altered immune regulation in Down syndrome to increased vulnerability to SARS-CoV-2–induced cardiac injury. More broadly, this work highlights the importance of considering genetic and chromosomal context in host–virus interactions and may inform the development of targeted therapeutic strategies aimed at modulating immune responses in high-risk populations.

## 4. Conclusion

In summary, this study demonstrates that cardiomyocytes derived from individuals with trisomy 21 exhibit increased susceptibility to SARS-CoV-2 infection and mount a markedly increased transcriptional response compared to euploid controls. While both groups activate a conserved antiviral program, trisomy 21 iPSC-CMs display heightened induction of inflammatory and immune signaling pathways alongside pronounced suppression of genes critical for cardiomyocyte structure, contractility, and metabolic function. Notably, these differences occur in the absence of significant variation in canonical viral entry factors, suggesting that post-entry mechanisms and dysregulated host responses are key drivers of the observed phenotype.

These findings may provide mechanistic insight into the increased cardiovascular risk and adverse clinical outcomes observed in individuals with Down syndrome during COVID. The exaggerated immune response identified here supports a model in which intrinsic hypersensitivity to inflammatory signaling contributes to both enhanced viral susceptibility and downstream cardiac dysfunction. More broadly, this work highlights the importance of host genetic context in shaping cellular responses to viral infection. Targeting dysregulated inflammatory pathways may represent a promising therapeutic strategy to mitigate cardiac injury in vulnerable populations such as those with trisomy 21.

## 5. Methods

### 5.1 Human Subjects

All human subject research was conducted under protocols approved by the Institutional Review Board at Nationwide Children’s Hospital. Written informed consent was obtained from all participants or their legal guardians prior to sample collection. Participants included female individuals with Down syndrome (trisomy 21) and their euploid, sex-matched biological relatives.

### 5.2 Generation of human iPSC lines

PBMCs were extracted from blood samples as previously described [10]. Briefly, 5 mL of whole blood were drawn from n=3 female subjects with trisomy 21 and n=3 of their healthy sex-matched biological parents. Blood was transferred to a cell separation tube (BD Vacutainer CPT tube, BD Biosciences) and centrifuged at 1,500 x g for 30 min at room temperature. PBMCs were then isolated from the middle layer of the fractionalized blood sample and transferred to a 15-mL conical tube. Cells were washed with 10 mL DPBS (Ca2+/Mg2+ free) by centrifugation at 300 x g for 15 min at room temperature. Cell pellets were resuspended in freezing media made of Knockout Serum Replacement (Gibco) plus 10 % DMSO and stored in liquid nitrogen until iPSC reprogramming.

Isolated PBMCs were cultured for 1 week at 37 °C, 5 % CO_2_ in StemPro-34 SFM medium (Thermo Fisher Scientific) supplemented with 1x GlutaMAX (Thermo Fisher Scientific), 20 ng/mL IL3 (PeproTech), 20 ng/mL IL6 (Gibco), 20 ng/mL EPO (Thermo Fisher Scientific), 100 ng/mL SCF (PeproTech), and 100 ng/mL FLT3 (Thermo Fisher Scientific). CytoTune™-iPS 2.0 Sendai Reprogramming Kit (Thermo Fisher Scientific) was used to transduce 5 x 10^5^ PBMCs. Cells were resuspended in supplemented StemPro34 SFM in Matrigel-coated plates (1:300 in DMEM-F12) for one week, after which cells were supplemented with complete E8 medium (Thermo Fisher Scientific). After two weeks, iPSC clones were picked, expanded, and preserved in liquid nitrogen.

### 5.3 iPSC maintenance and passaging

Human iPSCs were maintained in 6-well Matrigel-coated plates at 37 °C with 5 % CO_2_ in E8 medium until 90 % confluent. To passage, cells were washed with 3 mL DPBS and dissociated with 0.5 mM EDTA for 5–8 min. One well of iPSCs was replated using complete E8 medium supplemented with 2 µM ROCK inhibitor (Y-27632, Selleck Chemicals) at 1:6 ratio and allowed to recover overnight. The next day, medium was switched back to E8 medium.

### 5.4 Cardiomyocyte differentiation

Cardiac differentiation was performed using our established protocol [10]. Briefly, when iPSCs reached 90 % confluency, 8 µM CHIR99021 (Selleckchem) in basal differentiation media (RPMI1640 medium plus B27 minus insulin supplement) was introduced for 2 days at 37 °C with 5 % CO_2_. Media was then changed on day 2 to basal differentiation media and cultured overnight. On day 3, cells were then treated with 5 µM IWR-1 (Sigma) in basal differentiation media for 2 days, then exchanged with basal differentiation media for another 2 days. On day 7, cardiac maintenance media (RPMI1640 medium plus B27 supplement) was introduced every other day. Beating clusters were observed at days 8-12 of differentiation. Cultures were metabolically purified by removing glucose from the culture media for 4 days using RPMI1640 no glucose medium plus B27 supplement. Cells were then recovered in cardiac maintenance media for 4 days. To passage iPSC-CMs onto a fresh Matrigel-coated vessel, cells were washed with 3 mL of DPBS then incubated with 1mL of TrypLE Select Enzyme 10x (Thermo Fisher Scientific) for 5 min at 37 °C with 5 % CO_2_ to dissociate cells. The reaction was halted with addition of cardiac replating media (cardiac maintenance media plus 10 % KSR) and then centrifuged at 300 x g for 5 min. The supernatant was removed, and cells were resuspended and plated using cardiac replating media. Cells were then allowed to mature for 30 days by refreshing with cardiac maintenance media every other day.

### 5.5 Infection of human iPSC-CMs with SARS-CoV-2

SARS-CoV-2 USA-WA1/2020 infection experiments were performed in the Biosafety Level 3 facility at The Ohio State University. The SARS-CoV-2 USA-WA1/2020 natural isolate (BEI Cat: NR 52281) was used for all inoculations. Confluent DS and healthy control iPSC-CMs grown in 96-well plates were inoculated with 0, 50, 100, 200, and 400 PFU of the virus and incubated for 1 h at 37 °C. There were 5 replicates for each virus inoculation dose. The inoculum was removed after 1 h of viral absorption and replaced with 100 µL of cardiomyocyte maintenance medium. Infected cells were fixed with 4 % PFA on days 1 and 2.

### 5.6 RNA isolation and sequencing

Cells in 24-well plates were inoculated with 2000 PFU of SARS-CoV-2 and incubated for 1 h at 37 °C. Control wells were inoculated with media only. There were 4 replicates each for virus infection and control. The inoculum was removed after 1 h of viral absorption and cells were replaced with 500 µL of B27 medium. Cells were incubated for 2 days post infection and then lysed with 200 µL of Trizol. Lysates from quadruplet replicates were pooled, and the RNA was extracted using Zymo’s Direct-zol RNA Miniprep Plus kit. RNA was eluted in ultrapure water and sequenced using Novogene’s sequencing service.

### 5.7 Immunofluorescence

Differentiated cardiomyocytes were immunostained for cardiac-specific markers TNNT2 and alpha-actinin. SARS-CoV-2 was immunostained with nucleocapsid antibody (Sino-Biological, Cat: 40143-R001). Differentiated iPSC-CMs were fixed with 4 % paraformaldehyde solution (Electron Microscopy Sciences) for 15 min then permeabilized with 0.1 % Triton X-100 solution (Sigma) for 20 min at room temperature. Cells were then blocked with 0.2 % BSA (Sigma) in DPBS for 2 hours and then incubated overnight with primary antibodies (1:1000) at 4 °C. The next day, cells were incubated with secondary antibodies (1:2000) in 0.2 % BSA for 1 h at room temperature followed by counterstaining with DAPI (1:2000) for 5 min. Coverslips were mounted using SlowFade Gold Antifade (ThermoFisher Scientific) and imaged with a fluorescent microscope (Leica).

### 5.8 Bioinformatics analysis

Paired-end bulk RNA sequencing reads (150 bp) generated by Novogene were subjected to quality control, adapter trimming and low-quality base filtering using fastp (v0.23.2) with default parameters [16]. Clean reads were aligned to the human reference genome (GRCh38, Ensembl release 99) using HISAT2 (v2.2.1) [17] with default settings for paired-end alignment. The resulting SAM files were converted to BAM format, sorted by genomic coordinates and indexed using SAMtools (v1.17) [18]. Gene-level read counts were quantified from sorted BAM files using featureCounts from the Subread package (v2.0.1) [19], with the Ensembl GRCh38 release 99 GTF gene annotation. Only primary alignments were counted, and reads were assigned based on the gene name attribute. Genes with fewer than 20 total counts across all 24 samples were excluded from downstream analysis to remove lowly expressed transcripts.

Differential gene expression analysis was performed using DESeq2 (v1.44.0) [20] in R (v4.4.1). Raw count data and sample metadata were used to construct a DESeqDataSet object with the group variable (defined as the combination of diagnosis and infection status) as the design factor. Size factor normalization and dispersion estimation were performed using default DESeq2 parameters. Pairwise comparisons were conducted between infected (48 h post-SARS-CoV-2 infection) and baseline (uninfected) conditions within each diagnostic group: trisomy 21 COVID versus trisomy 21 control, and euploid COVID versus euploid control. Genes with a Benjamini-Hochberg adjusted P value less than 0.05 and an absolute log2 fold change of at least 2 were considered significantly differentially expressed. Variance-stabilizing transformation was applied to the normalized count data for principal component analysis and sample-level visualization.

Gene set enrichment analysis was performed using the fgsea package (v1.30.0) [21] in R with ranked gene lists ordered by the DESeq2 test statistic. Gene sets were obtained from the Molecular Signatures Database (MSigDB) [22], including Gene Ontology Biological Process (C5:GO:BP) and Reactome pathway (C2:CP:REACTOME) collections. Enrichment significance was assessed with 1,000 permutations, and pathways with an adjusted P value less than 0.05 were considered significantly enriched.

## Article Information

This study was conducted in accordance with protocols approved by the Institutional Review Board at Nationwide Children’s Hospital and written informed consent was obtained from all participants or their legal guardians. All experiments involving SARS-CoV-2 were performed in a certified biosafety level 3 facility at The Ohio State University in compliance with institutional and federal safety guidelines. The datasets generated and analyzed during the current study are available from the corresponding author upon reasonable request. Bulk RNA sequencing data have been deposited in the NCBI Gene Expression Omnibus (GEO) under accession number GSE332829.

## Acknowledgments

We gratefully acknowledge the clinical research team in The Heart Center and Dr. Kandamurugu Manickam in the Division of Genetic and Genomic Medicine at Nationwide Children’s Hospital for their role in recruiting patients with trisomy 21 and their euploid family members. We also thank Dr. Dennis Lewandowski for his assistance with manuscript editing.

